# Endoplasmic reticulum patterns insect cuticle nanostructure

**DOI:** 10.1101/2024.08.20.608717

**Authors:** Sachi Inagaki, Housei Wada, Takeshi Itabashi, Yuki Itakura, Reiko Nakagawa, Lin Chen, Kazuyoshi Murata, Atsuko H. Iwane, Shigeo Hayashi

## Abstract

Insect cuticles with nano-level structures exhibit functional surface properties such as structural color and superhydrophobicity. Despite the enormous influence the cuticle has had on biomimetic industrial applications, molecular and cellular mechanisms of cuticular extracellular matrix (ECM) assembly into nanoscale structures remain poorly understood in insects and other taxa. Ghiradella (1989) described how a crystallin-like lattice of endoplasmic reticulum (ER) prefigures the patterning of the porous cuticle of the butterfly wing scale with structural color^1^. Building on that insight, we show that the nanopore structure of the olfactory (olf) organs, which serve as molecular filters in *Drosophila,* is built through a novel process in which ER material is trafficked to the plasma membrane mediated by the autophagy pathway. The process is controlled by the insect-specific protein Gore-tex/Osiris23 (Gox)^2^, which is localized to the tubular ER of olf hair cells. Gox recruits Ref(2)P, the fly counterpart of mammalian p62/SQSTM1^3^, to initiate ER scission. The excised ER fragments are processed by autophagy to gain access to the plasma membrane and trigger membrane invagination, which plays a role in remodeling the cuticular envelope layer to the nanopore formation. This repurposing of the ER phagy for machinery to support the fabrication of nanoscale ECM by the Gox protein sheds light on the nanopatterning of insect cuticles and their genetic control.

## Main

Nanoscale modifications of the insect cuticle are the basis for various surface properties, such as efficient light transmittance and molecular filtering by sensory organs, and the selective light reflection that underlies intra- and inter-specific communication ^4–7^ These key structural innovations have enabled ancestral insects to explore and thrive in a wide range of terrestrial environments. The insect cuticle is a multi-layered structure of a great variety in thickness and chemical composition^8,9^. While the biochemical basis for the unique physical properties of cuticles has been investigated, how cuticular nanostructures are patterned at the cellular and genetic levels is poorly known.

The *Osiris (Osi)* gene family includes multiple candidate regulators of such nanoscale patterning. Molecular phylogeny has shown that these genes were acquired early in insect evolution, rapidly increasing in number and becoming durably conserved thereafter ^10,11^. In *Drosophila*, many of the 25 *Osi* genes are expressed in a variety of cuticle-secreting cells. Specific subsets of *Osi* genes are required for the patterning of the corneal nipple of the compound eye *(Osi9, Osi21)*, the tip pore of the taste hair *(Osi11)*, and nanopores of olf hairs (*gore-tex/Osi23*^2,12^). Nanopores are 30–50 nm pores on the surface of sensilla basiconica and sensilla trichordia, which function in the selective transport of air-born odorant molecules to olf neurons while excluding larger particles ^7,13^. Nanopore development starts as the wavy curving of the envelope layer, which tracks the curved and invaginated patterns of the plasma membrane in olf hair cells^2^. *gore-tex/Osi23 (gox)* is required for maintaining the curvature of the plasma membrane, which underlies nanopore formation and proper olfactory response^2^. Notably, ∼70% of the membranes showing endocytic structures are proximal to a downward curved envelope (Fig.1a), leading to the hypothesis that the plasma membrane serves as a template for curved envelope formation ^2^. Gox is known as a transmembrane protein localized to intra-membrane compartments, but its function in the plasma membrane and cuticular nanostructures is not well understood.

We now show the molecular pathway by which Gox exerts its patterning functions. Gox is localized to the endoplasmic reticulum (ER) and triggers ER phagy by recruitment of the autophagy mediator Ref(2)P^14^, a homolog of mammalian p62/SQSTM1^15^. The processed ER phagy product gains access to the plasma membrane and promotes plasma membrane invagination. In addition, Gox interacts with dynamin to maintain invagination and the shape of the olf organ. Thus, in insects, Osi-family protein Gox has exapted the conserved ER phagy pathway for nanoscale patterning of ECM.

### Gox promotes ER phagy during nanopore formation

Gox has been localized to intracellular vesicles that overlap endosomal markers ^2^. To determine the precise identity of the Gox-carrying organelle, we knocked the APEX2 tag^16^ into the N-terminus of the genomic *gox* gene. The tag, which did not affect Gox activity, allowed the labeling of the protein in transmission electron microscopy (TEM). APEX2 signal appeared in the luminal area of tubular ER structures (type I) surrounding the nucleus, and in the distal shaft region of the olf cells (sensilla basiconica of 3^rd^ antennal segment, An3, Fig. 1b, c). Additional label was found in electron-dense vesicle (type II), multi membrane structures (type IIIa), and vesicles with opaque staining (type IIIb, Fig. 1c). Type II vesicle had the highest level of Gox and was located most proximal to the plasma membrane (Fig. 1c, e). Type IIIa and type IIIb structures were sometimes located adjacent to invaginated membranes and curved envelope (Fig. 1c, d). A Type I - Type II - Type III transition was deduced from the assumption of a linear relationship (Fig. 1e, f). Consistently, fluorescently labeled HA-Gox overlapped partially with the tubular ER network in the central region of the olf cell, and densely localized underneath the plasma membrane, likely in Type II vesicle (Fig. 1g). This localization pattern indicates Gox is transported from ER into membrane-proximal structures.

**Figure 1a-c.**
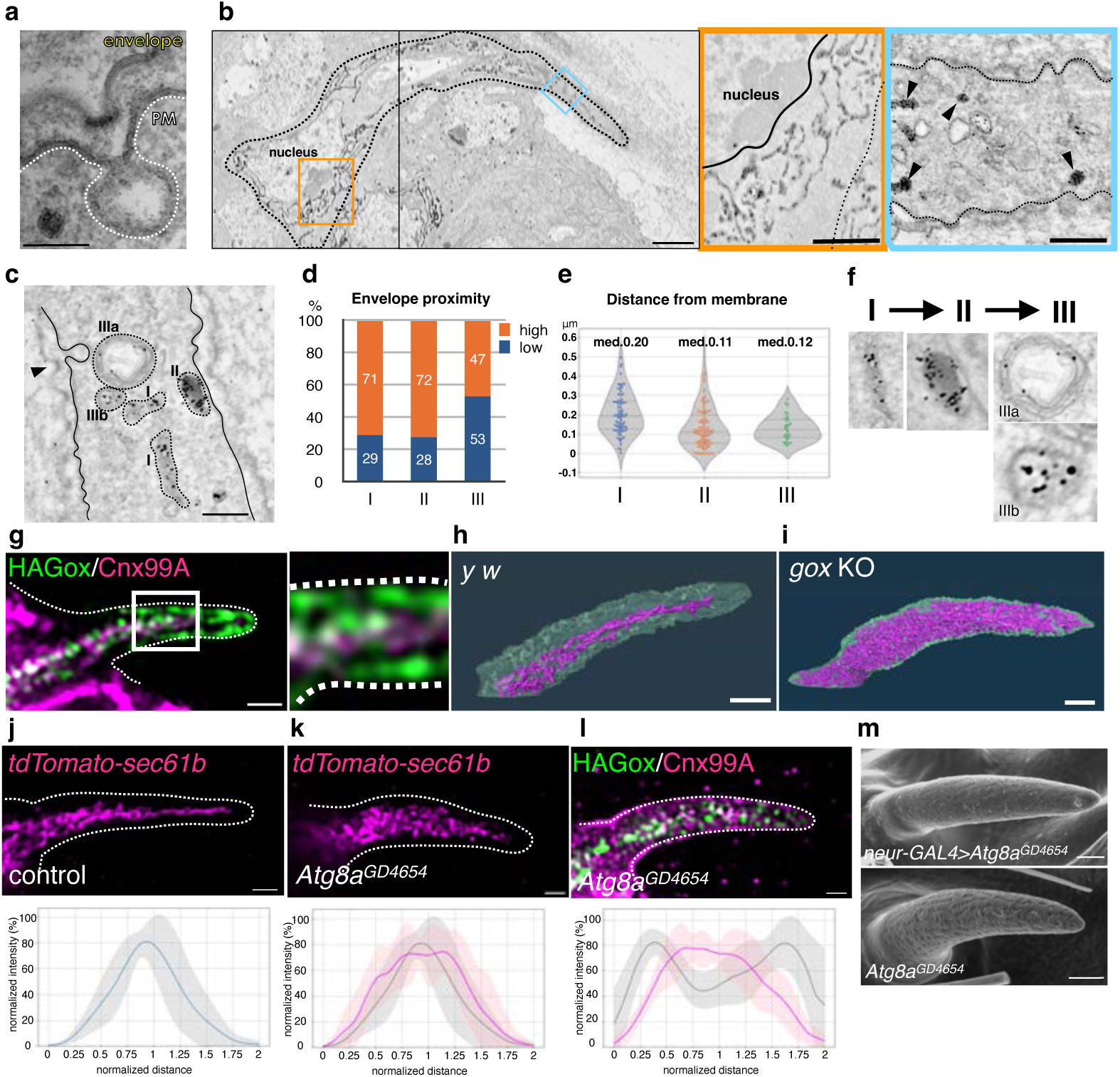
Transmission electron microscopy of the olf hair cell at 44h APF. **a,** The lowest point of the curved cuticular envelope is often associated with plasma membrane invagination. **b**, Localization of APEX2-Gox protein in tubular ER. **c**, Three types of Gox-containing organelles. I, ER. II, Electron-dense vesicle (arrowhead in **b**). IIIa, IIIb, Electron-lucent structures. IIIa is a multi-membrane structure. **d,** Proximity of Gox+ structures to the high or low point of the envelope. **e,** Distance from the membrane of the three types of Gox vesicles. Median distance is shown above each plot. **f**, Presumed order of Gox-vesicle conversion. **g,** Localization of Gox and ER marker Cnx. **h, i,** Distribution of ER in control (*y w*) and *gox* mutant. Segmented view of FIB-SEM stacks. **j-k,** ER distribution in control and ATG8a RNAi. **I,** Reduction of subcortical Gox localization in ATG8a RNAi. Graphs below show averaged line scans of ER or Gox along the line in **j** (control: gray, mutant: magenta, 5-7 hairs for each genotype). **m**, Adult olf bristle of neur>ATG8a RNAi lacking nanopores (top) and control (UAS-ATG8a RNAi only, bottom). Bar: 100nm (a), 2µm (b), 500 nm (enlarged in b), 200nm (c), 1µm (g-m).

In the olf cell, the ER formed an interconnected network extending along the cell major axis (Fig. 1h, Extended Data Fig. 1a). Endoplasmic reticulum network formation was poor in the non-olfactory hair (spinule, Extended Data Fig. 1c, d) In *gox* mutants, ER expanded to fill the entire cytoplasm (Fig. 1i, Extended Data Fig. 1b). Transmission electron microscopy of *gox* mutant olf hair cells revealed vacuolar structures with multi-layered membranes, similar to the ER whorls observed in ER stressed cells^17,18^ (Extended Data Fig1e).

The bulk amount of ER is negatively regulated by ER phagy under control of the autophagy pathway^15,19^. Knockdown of the core autophagy genes (ATG1, ATG2, ATG5, ATG8a, ATG18a) caused expansion of ER and concomitant reduction of Gox from the membrane-proximal region (Fig. 1j-l, Extended Data Fig. 2a, b). Mutant flies showed reduced nanopore formation (Fig. 1m, Extended Data Fig. 2c), suggesting that that ER phagy driven by Gox is essential to nanopore formation.

### Plasma membrane invagination is essential for the nanopore formation

To study the mechanics of plasma membrane patterning by Gox, we performed a 3D reconstruction of the plasma membrane using focused ion beam-scanning electron microscopy (FIB-SEM^20^). Olf hair cells showed a highly convoluted plasma membrane (plasma membrane convolution, PMC) on observation from the outside, whereas, the inside view showed numerous plasma membrane invaginations (PMI, 74 per hair, N=3, Fig. 2a, *y w*, Extended Data Fig. 3a). In *gox* mutants, the size of PMI was greatly reduced, while PMC was retained (Fig. 2a, middle row, Extended Data Fig. 3c). In contrast, spinules showed smooth membrane surfaces both outside and inside (Fig. 2a, bottom row, Extended Data Fig. 3b, d). These findings suggest that *gox* is involved in PMI formation, not PMC. Time course analysis showed the envelope in *gox* mutants is curved at 44h APF, gradually becoming flattened by 52h APF (Fig. 2b-d).

**Figure 2.**
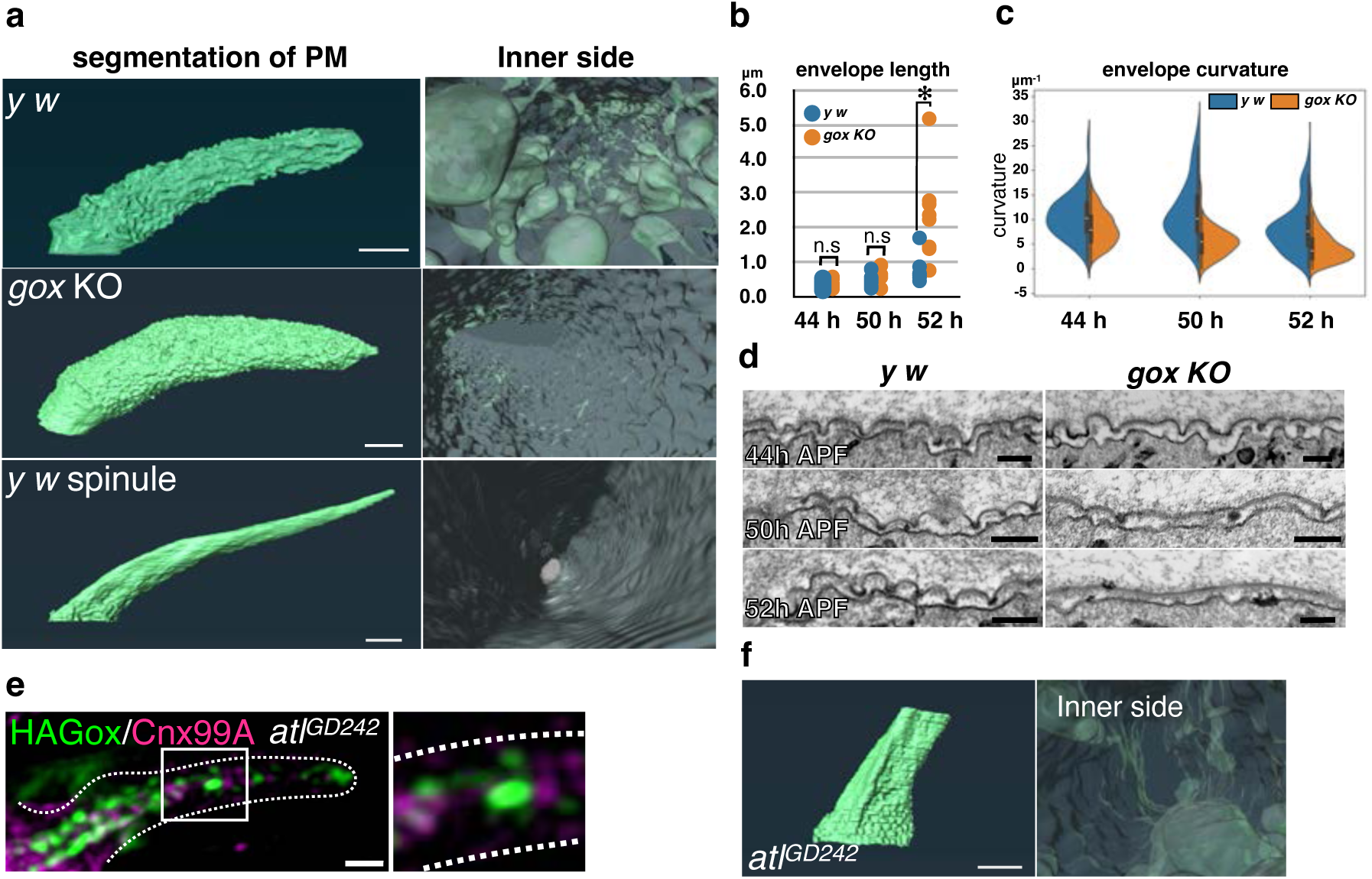
**a,** Plasma membrane structures of the olf and spinule obtained by FIB-SEM. Outer view (left), inner view (right). **b**, The length of the envelope measured between low points. Significance was determined by Mann-Whiteny U test. **: p<0.01, n.s.: not significant. **c**, Quantification of envelope curvature (μm^-1^). **d,** Longitudinal TEM views of the plasma membrane and envelope. **e**, ER, and Gox in *atl* RNAi. **f**, FIB-SEM plasma membrane views of *atl* RNAi hair cell.

To address the role of ER, we knocked down the ER fusion protein Atlastin (Atl)^21^. This treatment caused ER fragmentation and the appearance of ER whorl (Extended Data Figure 4e, f), reduced cortical Gox distribution (Fig. 2e), the loss of PMI and PMC (Fig. 2f), and flattening of the envelope at 44h APF (Extended Data Figure 4e). In the adult maxillary palp, *atl* knockdown caused near-complete loss of olf organs, while the number of similarly treated mechanosensory bristles was unchanged (Extended Data Figure 4a-c).

These results indicate that the ER network is required for PMC and PMI formation. Moreover, restricting the amount of ER by Gox-mediated ER phagy is essential for PMI formation and maintenance of envelope curvature, itself required for nanopore formation.

### Gox recruits Ref(2)P/p62 to the plasma membrane

To identify Gox effector molecules, we immunoprecipitated Gox-associated proteins from S2 cells and analyzed them by mass spectroscopy. Molecules identified included proteins involved in ER quality control (Calnexin, Sec61alpha) and molecular chaperons (Hsc70), as well as molecules involved in membrane dynamics (Fig. 3a, Supplementary Table 2). Representative molecules (Sponge, Rac1 GEF; TER94, ATPase for ER membrane budding; CG13887, ER to Golgi transport) were expressed in S2 cells and were confirmed to be ER-localized (Fig. 3b). RNAi knockdown of *spg, TER94, CG13887, Kr-h2* and *Ref(2)P* caused reduction of nanopores in the olf bristle (Fig. 3c).

**Figure 3.**
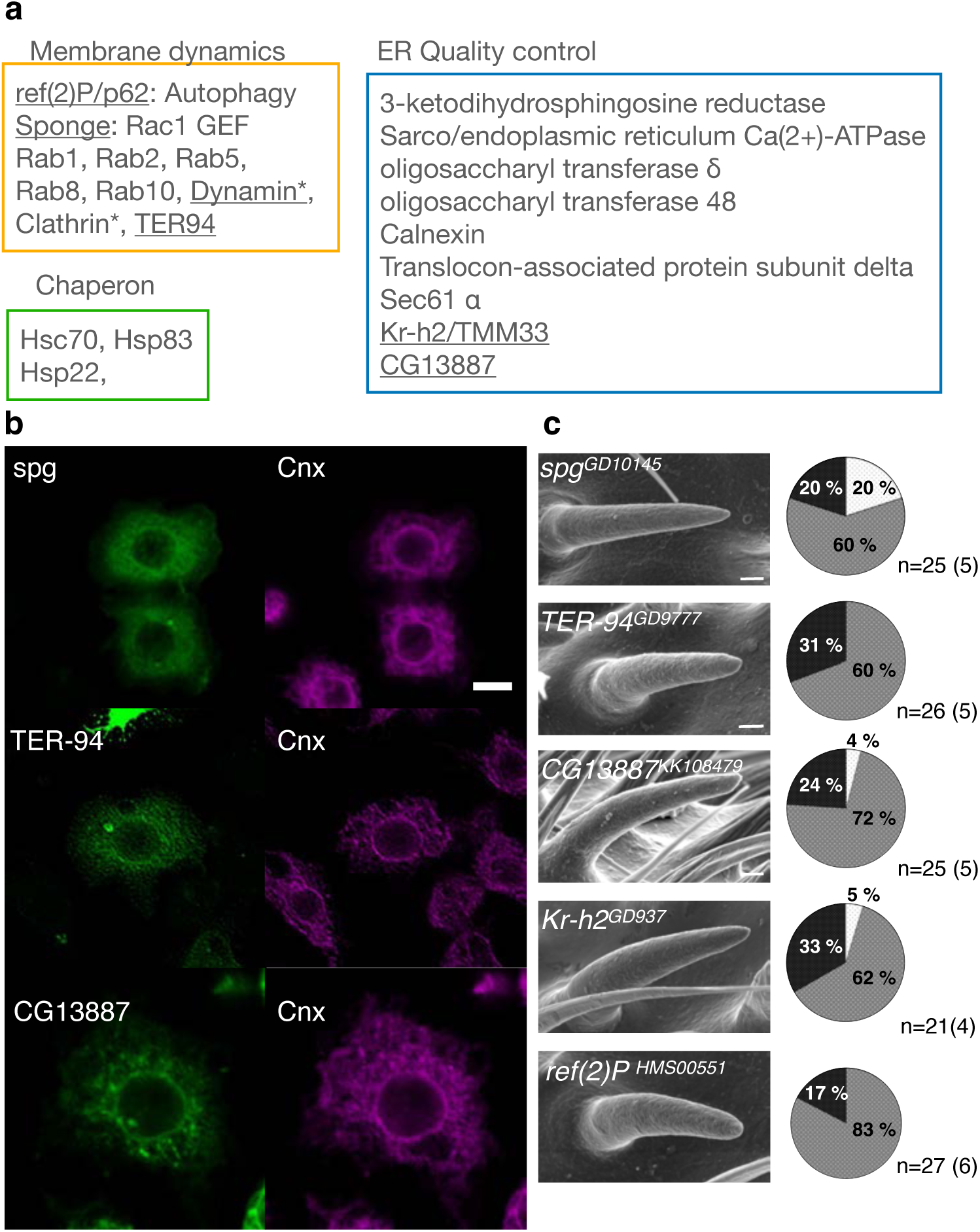
**a,** Mass spectrometric identification of Gox interacting proteins in S2 cells expressing Gox. Asterisk (*) indicates proteins identified from cells co-expressing Gox and Ref(2)P. **b,** ER localization of Gox interacting proteins Spg, TER-94, CG13887. Image of TER-94 was processed by image deconvolution. **c,** RNAi phenotypes of Gox interacting proteins/genes. SEM image of olf bristles. Left: SEM image of olf bristles showing strong loss of nanopore phenotype. Right: percentage of olf bristles classified as normal (Light gray), partial (gray) and strong (black) class of nanopore phenotype.

We focused on Ref(2)P, a homolog of mammalian p62/SQSTM that functions as an adaptor molecule linking the autophagy molecule LC3 and protein ubiquitination^14,15^. While Ref(2)P was ubiquitously localized as cytoplasmic puncta in *Drosophila* tissues^14^, it was highly enriched in the membrane-proximal region of the olf hair cell, and a subset of it overlapped with HA-Gox (Fig. 4a,b). *gox* mutation reduced the membrane-proximal Ref(2)P (Fig. 4b lower panel), and Ref(2)P knockdown reduced membrane-proximal Gox localization (Fig. 4c). These results suggest that Gox and Ref(2) interact to co-localize to the plasma membrane.

**Figure 4.**
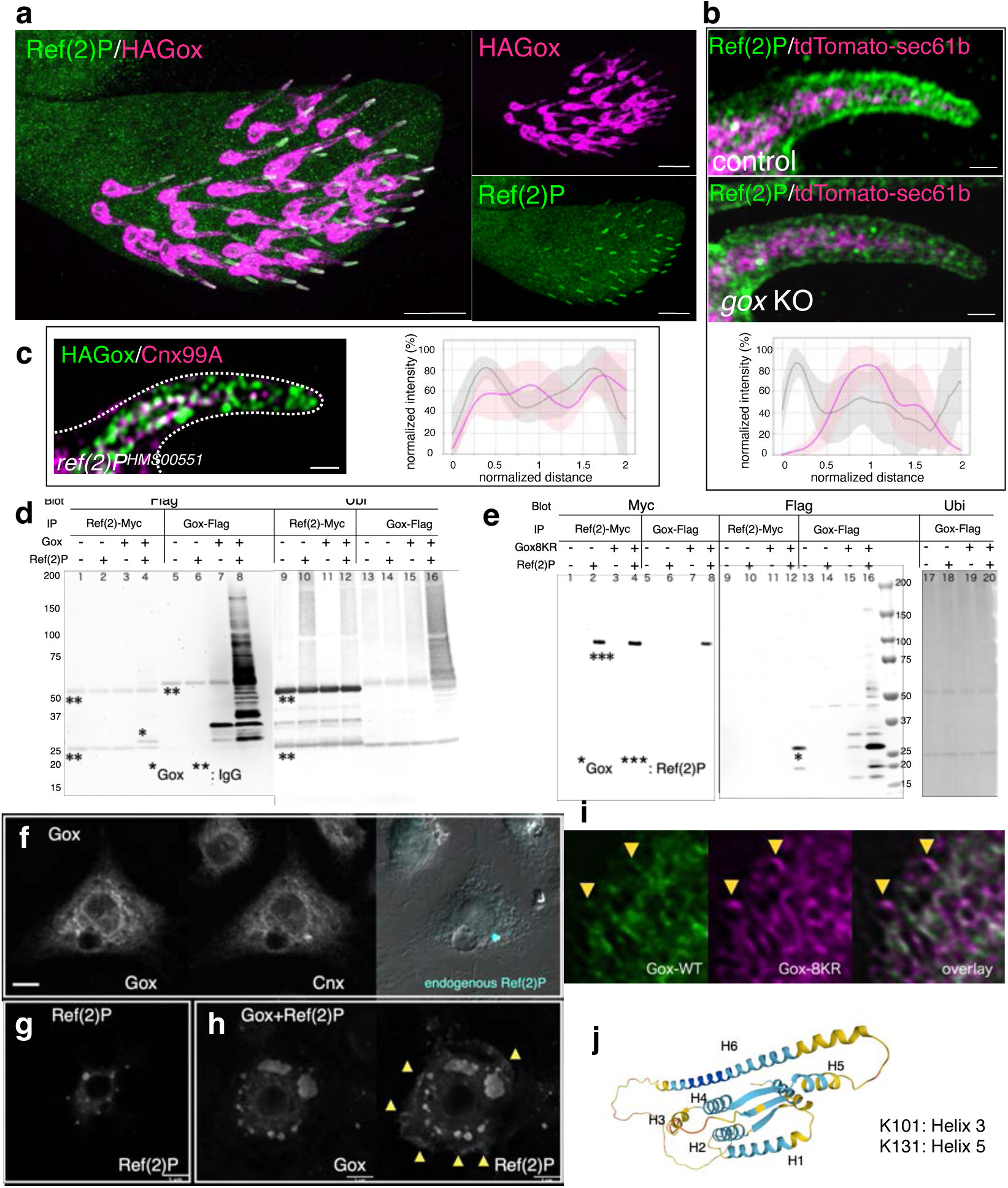
The role of Ref(2)P in Gox localization and ubiquitination. **a,** Ref(2)P is specifically enriched the shaft region of olf hair cells expressing HAGox. Maxillary palp 42h APF. **b,** Subcortical localization of Ref(2) in control and *gox* mutant olf hair cells, and their quantification**. c,** Subcortical Gox localization was lost in Ref(2)P RNAi. Right graph: quantification. **d,** Gox-Flag and Ref(2)P-Myc were coexpressed and immunoprecipitated with the tag antibodies. Immunoprecipitate of Ref(2)P contained Gox (lane 4). The increased amount of high-molecular-weight forms of Gox was induced by Ref(2)P (lane 8). Note that cross-reactivity of HRP-conjugated antibodies to the IgG (marked with **) used for immunoprecipitation. **e,** Properties of Gox8KR. Gox8KR and Ref(2)P were co-immunoprecipitated with each other (lane 8, 12). Although Ref(2) caused an increase in Gox amount and molecular mass (lane 16), no specific increase in ubiquitination level was observed (lane 20). **f,** In S2 cells, Gox is localized to ER labeled with Calnexin. Endogenous Ref(2) occasionally formed condensates associated with ER. **g,** Overexpressed Ref(2)P formed multiple condensates. **h,** Coexpressed Gox and Ref(2)P colocalized in condensate. Note the increase of Ref(2)P at the cell margin (arrowhead). Bar: 5µm. **i,** Differential ER localization of wild type Gox and Gox8KR. **j,** alpha-fold 2 model of Gox/Osi23 and the location of the two mapped ubiquitinated lysine.

Gox expressed in cultured S2 cells was detected as multiple bands of relative molecular mass of 25-30 kd (Fig. 4d, lane 7). Co-expression of Ref(2)P increased the amount of Gox and its molecular size up to 150kd that reacted with anti-polyubiquitin antibody (Fig. 4d, lane 8, 16). Although mammalian p62 is known to bind ubiquitin, Gox bound to Ref(2)P was exclusively the native form of 28.6 kd with no ubiquitin (Fig. 4d, lane 4). In S2 cells, endogenous Ref(2)P formed condensates and Gox was present in ER (Fig. 4f). Ref(2)P condensate was increased upon its overexpression (Fig. 4g). Co-expression of Gox and Ref(2)P caused the inclusion of Gox in the Ref(2)P condensates, and a subset of Ref(2)P was translocated to the peripheral membrane (Fig. 4h, arrowhead). We identified two ubiquitination sites, K101 and K131, by mass spectrometry (Fig. 4j, Extended Data Fig. 5c, d). Substitution of the last eight, or all 11 lysines of Gox to arginine (Gox8KR, Gox11KR, including K101 and K131) eliminated ubiquitination, but preserved its binding to Ref(2)P (Fig. 4e, Extended Data Fig. 5b). Substitution of the last four lysines, leaving K101 and K131 intact (Gox4KR), resulted in identical binding function to that of wild type Gox (Extended Data Fig. 5a). Gox8KR was localized to ER, but a subset of it was segregated from wild type Gox (Fig. 4g). These data indicate that ubiquitinated Gox is localized to ER, and its deubiquitination permits association with Ref(2)P.

### Plasma membrane invagination is the site of ER-membrane contact

Plasma membrane invagination resembles the clathrin-coated pit that forms during endocytosis, a rapid process completed in ∼90 sec by dynamin-catalyzed scission^22^. To capture the interaction of PMI and ER in the snapshot images of EM, the PMI scission reaction was temporally arrested by reducing the activity of dynamin using the *shibire^ts^*^2^ (*shi^ts^*^2^) mutation^23^. A temperature shift was applied at the time equivalent of APF 36h to 37h at 25°C, following the protocol optimized to maximize the effect on olf hair formation while avoiding developmental arrest (Fig. 5a). This transient inactivation of dynamin was sufficient for causing prominent effects in the olf hair cells.

**Figure 5.**
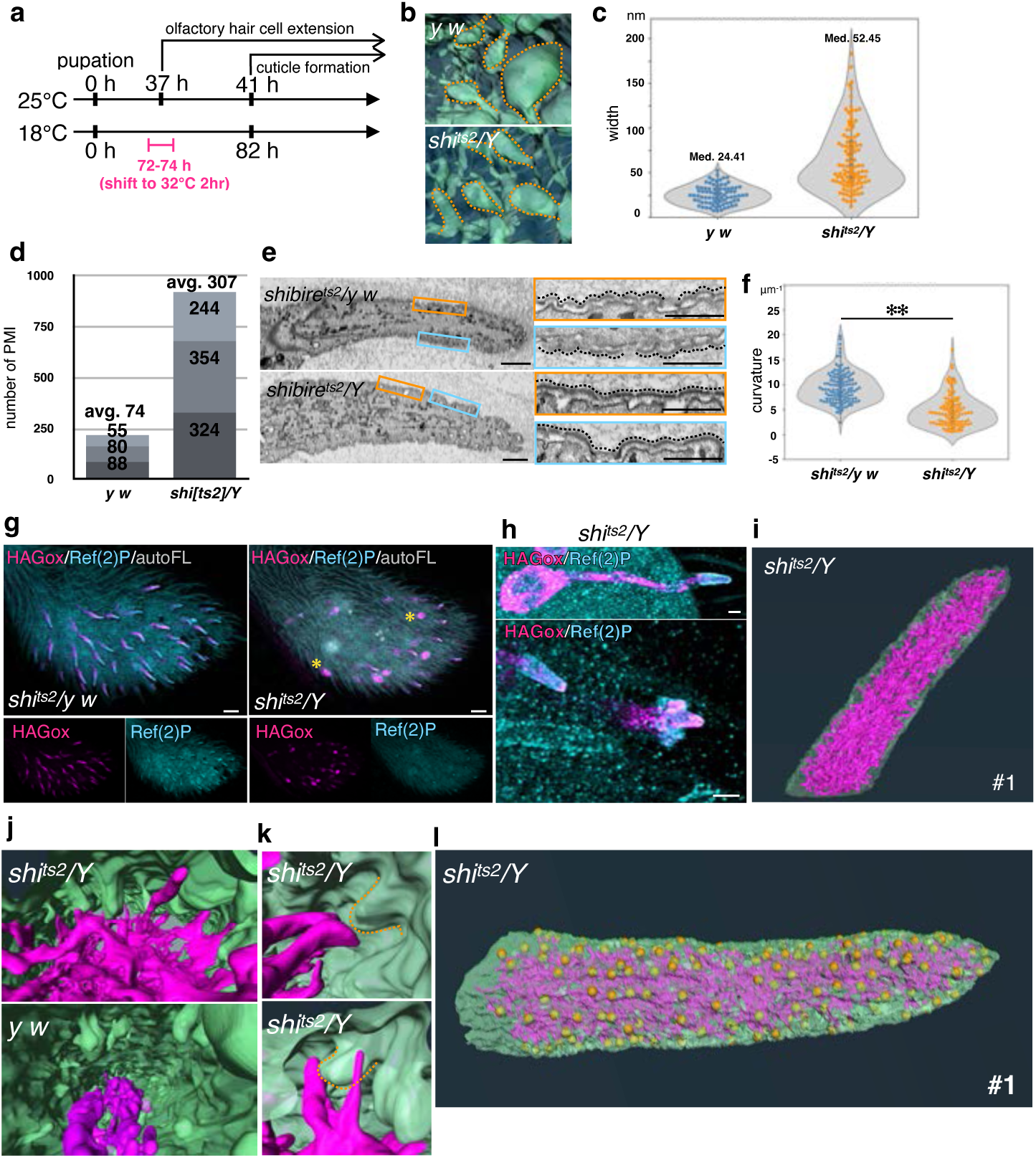
Dynamin-Gox interaction. **a,** Temperature-shift protocol for inactivating dynamin in *shi^ts^*^2^ mutants. **b**, Plasma membrane invagination (PMI) in control and *shi* mutants. **c**, PMI neck width measurement. **d**, Number of PMI in 3 hair cells of each genotype. **e,** TEM views of *shi^ts^*^2^/*y w* (control) and *shi^ts^*^2^/*Y* (mutant). **f,** Quantification of envelope curvature. Significance was determined by Mann-Whitney U test **: p<0.01. **g**, Maxillary palp of *shi^ts^*^2^/*y w* (control) and *shi^ts^*^2^/*Y* (mutant) stained for expression of HA-Gox and Ref(2)P. Asterisk indicates completely invaginated olf hair cells. **h**, Enlarged views of *shi^ts^*^2^/*Y* olf hair cells, showing normal (top and lower-left), and partially invaginated (lower right) phenotypes. **i.** FIBSEM view of plasma membrane and ER of *shi^ts^*^2^/*Y* olf #1. **j,** Relationship of ER and plasma membrane in *shi^ts^*^2^/*Y* and *y w*. **k,** ER-plasma membrane contact interface in *shi^ts^*^2^/*Y*. **l,** Distribution of ER-plasma membrane contact site (yellow dot) in *shi^ts^*^2^/*Y* olf. Bar: 1µm (**e**, left), 500nm (**e**, right), 10µm (**g**), 1µm (**h**).

Measurement of the 3D views of control and fully elongated *shi^ts^*^2^ olf hair cells at 44h APF (at 25°C) revealed the increase in the neck width and depth of PMI (Fig. 5b, c, Extended Data Fig. 6a), and the number of PMI (Fig. 5d). TEM views showed a reduction of envelope curvature in elongated olf hair cells (Fig. 5e, f). The cortical localization of HAGox and Ref(2) were normal (Fig.5 g, h). The ER was markedly over-expanded in olf hair cells, and numerous tubular ERs were extended toward the plasma membrane (Fig.5i). In magnified views, frequent association of PMI and PMC with extended ER was observed (Fig. 5j, k). The results demonstrated that PMI is undergoing frequent dynamin-driven scission coupled to Gox-dependent ER-phagy. *shi^ts^*^2^ caused the arrest of this dynamic process, suggesting a mechanistic link between ER and PMI. The ER-plasma membrane contact sites mapped on the surface of *shi^ts^*^2^ olf hair cells showed a scattered appearance resembling the nanopore pattern in wild-type olf bristles (Fig. 5l, Extended Data Fig. 6d).

### Plasma membrane invagination links hair cell shape to ECM nanopatterns

The emerged flies of *shi^ts^*^2^ showed a significant reduction in olf bristles, whereas the numbers and shapes of mech bristles and spinules were unaffected (Extended Data Fig. 6b). High-magnification imaging of the olf bristles showed variable phenotypes of full elongation, malformation, and complete loss, leaving a hole in the cuticle (Extended Data Fig. 6c). At APF44H, olf hair cells showed various morphologies including normal elongation, cupshaped indentation, and a total internalization subjacent to the epidermis (Fig. 6g asterisk, h). Olf hair cells expressing the dominant negative *shi^ts^*^1^ construct showed a similar internalized phenotype (Extended Data Fig. 6e). By labeling the olf ECM marker Tyn, we visualized hair cell-shaped ECM remnants associated with totally invaginated olf hair cells (Extended Data Fig. 6e), suggesting that dynamin deficient olf hair cells elongated and then retracted inward at 44h APF, leaving empty ECM outside.

Dynamin was found among the Gox-associated proteins (Fig. 3a) and 3D analysis of the plasma membrane revealed *gox* and *shi^ts^*^2^ showed opposite phenotypes in the PMI volume and depth (Extended Data Fig. 6a). We next asked whether the two genes genetically interact. The combination of *gox*-RNAi to *shi^ts^*^2^ caused significant suppression of the loss-of-olf and epidermal hole phenotypes of *shi^ts^*^2^ in the adult (Extended Data 6f, g), suggesting that PMI is the point of Gox function that links the dynamics of the ER and the plasma membrane.

## Discussion

Cuticles are secreted from the membrane in a variety of forms. Flat envelopes are produced from epidermal cells with membrane protrusions (plasma membrane plaque^24^), and hair cells with flat membranes rich in subcortical actin filaments (mechanosensory bristle^25^, spinule, this study) secrete flat envelopes. The olf hair cells are unique in that the entire plasma membrane is convoluted (PMC) and occasionally invaginated (PMI) and produces a wavy envelope and nanopores. The full ER function is required for shaping these plasma membrane structures, since PMI and PMC are lost in *atl* RNAi, and the envelope curvature did not appear at 44h APF in mutants. *gox* specifically affects PMI and the maintenance of envelope curvature. Our results suggest two-step regulation of envelope curvature by ER. The initial curvature at 44h APF is induced by the full activity of the tubular ER network, possibly the envelope following the convoluted plasma membrane (step 1). Gox maintains this curvature by promoting PMI and ER interaction, and PMI helps to maintain and further develop the curved envelope (step 2). The ER network contributes to step 1 by providing excess membrane to form PMC. Gox contributes to step 2 by promoting PMI formation to maintain the envelope curvature.

Gox is present in the ER membrane and transported to the shaft part of the olf hair cells, where it converts a fraction of ER into type II vesicles that are further changed to type III vesicles with autophagosome-like morphology. Ubiquitination of Gox may contribute to a change in ER membrane curvature and scission, as reported for the ER membrane protein FAM134B^26–28^. Deubiquitylation of Gox after scission may allow the interaction with Ref(2)P to initiate autophagic processing of the ER membrane. Autophagic ER membrane gains access to the plasma membrane to initiate PMI formation, possibly mediated by the interaction of Gox with dynamin and clathrin.

Inhibition of dynamin simultaneously arrested endocytosis and ER phagy, suggesting the two processes requiring Gox are functionally coupled at PMI. This coupling allows access to the plasma membrane by the ER, enabling the transfer of lipids for expanding the plasma membrane surface that convolutes in the limited space covered by the apical ECM. The ER may also provide lipids and secreted materials through PMI to the prospective pore region of the curved envelope, which is detergent-sensitive^2^. The space between the envelope and plasma membrane is filled with apical ECM that includes the zona pellucida-like domain protein Dusky-like (Itakura et al., Submitted). Dynamin was identified as a microtubule-associated protein^29^, and prolonged inactivation of temperature-sensitive dynamin caused microtubule bundling^30^. Coupling of microtubule bundles to dynamin may apply a pulling forth to PMI, and the force is likely to be transmitted to the envelope to mechanically promote envelope curving.

The similarity of the nanopore distribution and the pattern of ER-plasma membrane contact in *shi^ts^*^2^ olf hair cells supports a model in which the ER network serves as a template for nanopore patterning, as previously suggested in butterfly scale cells, where complex invagination of the plasma membrane associated with ER precedes the formation of cuticular photonic nanocrystals^1,31–33^. The template function of ER in specifying plasmodesmata, the nanoscale (50 nm diameter) intercellular channel across the plant cell wall is documented^34,35^. Gox/Osi23, an insect-specific innovation, allowed the ER to gain its novel function in cuticle patterning through coupling to the autophagy pathway. Future investigation into the roles of *Osiris* genes in other nanostructures in *Drosophila* (e.g., corneal nipples and tip pore) and structural colored scales in the butterfly and beetle will open rich new possibilities in the manipulation and design of insect cuticles with novel or modified surface functions.

## Methods

### Fly stocks and husbandry

The flies were maintained on standard corn meal-yeast food at 25°C unless otherwise noted. Fly pupae at the white prepupal stage were picked up and staged. Fly stocks are listed in Supplementary Table S1.

### Pupal tissue preparation

For fluorescent microscopy, olf hair cells of maxillary palp were analyzed. For FIB-SEM, TEM and APEX2 analyses, the dorsal-medial region of the third antennal segment rich in olf organs was chosen for analysis. Gox/Osi23 is expressed in both tissues and is required for olf nanopore formation^2,12^.

### Temperature shift experiment

The temporal inactivation of Dynamin using *shi^ts^*^2^ mutation was designed according to the developmental time course in different temperatures^36^. Virgin females of *shi^ts^*^2^ were crossed to *y*^1^ *w*^1118^ males, and the progenies were cultured at 18°C before staging at the white prepupal stage. Pupae at 72h APF (equivalent to 36h APF at 25°C) were heat-treated at 32°C for 2 hours in 1.7 ml microfuge tube placed in a block incubator and then returned 18°C until eclosion. Males (*shi^ts^*^2^*/Y,* mutants) and females (*shi^ts^*^2^/*+,* heterozygote control) were analyzed by FE-SEM. For genetic interaction assay with *gox* RNAi, male *gox ^kk^*^103932^; *neur-GAL4/TM6c SbDfd-GMR-YFP* strain was crossed to virgin *shi^ts^*^2^ females.

### APEX2 tagged-gox knock-in vector construction

APEX2 was targeted to the C-terminus of the Gox signal peptide so that APEX2 is retained to the N terminus of the mature Gox protein after signal peptide removal. Oligonucleotides used in this experiment are listed in Supplementary Table 2. The knock-in strain with APEX2 tagged-gox was constructed by following the scarless gene editing protocol from the laboratory of Kate O’Connor-Giles (https://flycrispr.org/scarless-gene-editing/). To build the donor plasmid, 1 kb each of the 5’ and 3’ homology arms flanking the target site were amplified using KOD-One (Toyobo) from the *gox* genomic rescue construct^2^. The APEX2 sequence amplified from pAPEX2-Dyl, and the 3xP3-DsRed cassette with PBac transposon ends were amplified. The sequence of gRNA target sites in the homology arms were mutated using the primer set SI7-SI8 and SI9-SI10 to prevent cleavage of the donor vector. The four fragments were ligated using the In-Fusion HD Cloning kit (Takara) into the pBS SKII vector cut with EcoRI (TAKARA). The plasmid sequence was confirmed by DNA sequencing.

Three gRNAs to the sequences around 300 bps upstream and downstream of the target site were designed using Target Finder (http://targetfinder.flycrispr.neuro.brown.edu) and CRISPR efficiently Predictor (https://www.flyrnai.org/evaluateCrispr/input), and were inserted into the BbsI-HF site of pCFD3-dU6 plasmid (Addgene #49410).

### Genome Editing

A mixture of the three gRNA vector plasmids (50 ng/μl each) and the donor plasmid (150 ng/μl) was injected into the *y*^2^ *cho*^2^ *v1 P{nos-Cas9, y+, v+}1A/Cyo* strain. Transformed flies were identified based on the red fluorescence in the eye, and balanced strains were established. Those flies were crossed with *w*^1118^*; CyO, P{Tub-PBac/T}2/wg^Sp-^*^1^*; l(3)/TM6B, Tb1* to excise the 3xP3-DsRed element and fluorescence-negative individuals were selected. Correctly edited strains were identified by PCR and DNA sequencing.

### Detection of APEX2 by electron microscopy

44-45 h APF pupae were removed from the pupal case and cut open in the abdomen, and were transferred to fixation buffer (2.5% glutaraldehyde, 2% formaldehyde, 0.1 M sodium cacodylate buffer, pH 7.4) on ice and were stored at 4°C for overnight. The tissues were washed three times for 10 min each in 0.1 M cacodylate buffer. Pupal cuticles and the body were removed from the head tissues and were washed with 0.1 M cacodylate buffer 3 times for 10 min.

DAB treatment and silver-gold enhancement were performed using a modification of the published method^37^. Fixed heads were incubated in 200 mM Glycine in 0.1M cacodylate buffer for 5 min on ice, rinsed with 0.1 M cacodylate buffer for 10 min. DAB solution was prepared by mixing 2 mg/ml DAB in H_2_O (prepared from DAB tablet, Sigma) with an equal volume of 0.1 M sodium cacodylate buffer on ice and filtering through a 0.22 μm syringe filter. Fly heads were incubated in 500 µl of DAB solution on ice for 30 min. The solution was replaced with DAB solution containing 10 mM H_2_O_2_ and incubation was continued on ice for 30 min to 1 hour. After the DAB reaction, samples were washed three times with 0.1 M sodium cacodylate for 10 min, with H_2_O 4 times for 15 min each, and 1% BSA in 20 mM glycine for 20 min, on ice. Next, the samples were incubated in 1% BSA and 20 mM glycine for 20 min at room temperature and prewarmed at 60°C for 10 min, followed by incubation in prewarmed silver enhancement solution (20:1:2 mixture of 3% hexamethylenetetramine, 5% silver nitrate, and 2.5% disodium tetraborate) for 15 min at 60°C for 15 min. Samples were washed with H_2_O 3 times for 5 min at room temperature, incubated with 0.05% tetrachlorogold (III) acid trihydrate in H_2_O for 5 min at room temperature, washed in H_2_O, and incubated with 2.5% sodium thiosulphate for 4 min at room temperature. Samples were washed H_2_O, incubate with 2.5% sodium thiosulphate for 4 min at room temperature, and then washed with H_2_O for 3 times. The samples were post-fixed with 1% osumium tetroxide and 1% (wt/vol) potassium ferrocyanide in H_2_O. After washing three times for 5 min each in H_2_O, samples were immersed in 1% uranyl acetate at 4°C for overnight. The stained tissues were washed three times for 5 min each in water, the tissues were subsequently dehydrated in a graded ethanol series. Following embedding and sectioning procedures were the same as for the FIB-SEM and the TEM procedure described below.

### 3D cell imaging by focused ion beam/scanning electron microscopy (FIB-SEM)

Pupal head fixation followed the procedure for APEX2 staining up to the removal of pupal cuticles. Samples were washed three times for 10 min each in 0.1 M cacodylate buffer, and post-fixed in 2% OsO_4_, 1.5% (wt/vol) potassium ferrocyanide, 2 mM CaCl_2_, in 0.15 M sodium cacodylate buffer (pH 7.4) for 2 h on ice in the dark. The fixed tissues were then washed three times for 10 min each in water and immersed in 1% thiocarbohydrazide (TCH) for 1 hr at 60°C. The tissues were then washed five times for 5 min each in water at room temperature. and fixed again in 2% OsO_4_ for 1 h on ice. The fixed tissues were washed three times for 5 min each in water, and stained *en bloc* in 1% uranyl acetate in H_2_O overnight at 4°C. The stained tissues were washed three times for 5 min each in water, and the tissue was stained with lead aspartate solution at 60°C for 1 h. The tissues were washed three times for 5 min each in water at room temperature, dehydrated in a graded ethanol series (30, 50, 70, 80, 85, 95, and 99.5%) each for 5 min on ice, and transferred to 100% ethanol for two 15-min on ice. After dehydration, the tissues were incubated in icecold 100% acetone for 5 min and incubated for 5 min with 100 % acetone at room temperature. For resin substitution, samples were immersed in a 3:1 mixture of 100 % acetone and Resin (8.8g Epon812, 2.7g DDSA, 0.23 ml DMP-30) at room temperature overnight, 1:1 mixture of aceton/Resin for 6 hrs, and in 1:3 acetone/Regin for overnight. Then samples were incubated in 100% resin at room temperature overnight. The Resin was replaced by fresh one and was polymerized by sequential incubation at 60°C 12 h, 45°C 12 h, 65°C 48 h, and 70°C 24 h.

After complete polymerization, the excess resin was trimmed, and the tissue block was mounted on an aluminum pin with electrically conductive glue, with the back side of the head on the top. Using an ultramicrotome (EM UC7 microscope, Leica), the block was cut from the back to front direction until the antennae were exposed, and coated with osmium (Tennant 20, Meiwafocis). Serial FE-SEM imaging of the block surface cut with a focused ion beam was performed (Helios G4 UC and Aquilos2, Thermo Fischer Scientific). Typically, 4 x 4 x 10 nm voxel (x-y-z) images (5000x, ∼1000 sections, 1.5kV, 0.1nA) of 10 x 20 µm area were obtained from the inside to the outside direction at the medial-dorsal region of the third antennal segment rich in sensilla basiconica.

### Data segmentation and analysis

Automatic alignment of image stacks and signal normalization between slices, 3D reconstruction, and manual segmentation of the image stacks were performed using the Amira software Version 2020.2 (Thermo Fisher Scientific).

### Transmission electron microscopy (TEM)

Sample fixation and embedding were performed according to the protocol of FIB-SEM sample preparation. After trimming the embedded tissue, ultrathin sections (approximately 50 nm) were cut and mounted on 200 mesh copper grids. Specimens were observed using a JEM-1400Plus transmission electron microscope (JEOL) at 100 kV accelerating voltage.

### Field Emission Scanning electron microscopy (FE-SEM)

The adult heads were dissected in PBS and then rinsed with 0.1 M cacodylate buffer three times for 5 min each, and incubated in fixation buffer 1 (2% paraformaldehyde, 2.5% glutaraldehyde, 0.1 M cacodylate buffer) at 4°C overnight. The samples were rinsed with 0.1 M cacodylate buffer three times at room temperature, for 5 min each. Then, the samples were incubated in fixation buffer 2 (1% OsO_4_, 0.1 M cacodylate buffer) on ice for 120 min in a light-shielded condition. Samples were further rinsed in water three times on ice with the light-shielded condition and subsequently dehydrated in a gradient of ethanol concentration, from 25%, 50%, 75%, 80%, 90%, 95%, 99.5%, and 100% for 10 minutes each at room temperature. The final 100% ethanol was dehydrated by the addition of a molecular sieve (Nacalai Tesque). The samples were dried by storing in the desiccator for 3-4 days. After dehydration, the heads were mounted on double-sided carbon tape on a brass pedestal and coated with OsO_4_ at approximately 15 nm thickness using an osmium coater (Tennant 20, Meiwafosis Co., Ltd.). The samples were observed with the field emission scanning electron microscope (JSM-IT700HR, JEOL).

### Fluorescence microscopy

Super-resolution images of the olfactory sensilla and spinules on the maxillary palp were acquired by the confocal microscopes equipped with Plan-Apochromat 63x/1.4NA oil immersion objective lenses and the Airyscan detectors (Zeiss LSM 880 and LSM980). Images were reconstructed by Airyscan processing with the 3D or 2D auto setting of the Zen software (Carl Zeiss) and analyzed with imageJ/Fiji^38,39^.

For sample preparation, 44h APF pupae were dissected out from the pupal case and fixed with 4% paraformaldehyde and 0.3% Triton X-100 in PBS for overnight at 4°C. Pupal cuticles were removed from the head with sharp forceps (Inox #5). Pupal heads were washed three times for 10 min with PBST (0.3% Triton X-100 in PBS) at room temperature. Samples were blocked with PBSBT (0.3% Triton X-100 and 0.1 % BSA in PBS) for 20 min at room temperature and incubated with 1^st^ antibody solution in PBSBT overnight at 4°C. Samples were washed three times with PBST and once with PBSBT, each for 10 min, and incubated with 2nd antibody in PBSBT at room temperature for 2 hr. After washing three times for 10 min with PBST, the heads were incubated in the mounting solution (SlowFade Diamond Antifade Mountant ThermoFisher, S36972). The maxillary palp was detached from the head and mounted on a slide glass.

The following antibodies and staining reagents were used: rat anti-HA (1:500; Roche, 3F10), mouse anti-Calnexin99A (1:10, DSHB, Cnx99A 6-2-1) and rabbit-anti RFP(1:200, BD Bioscience, and 1:200, MBL), rabbit anti-Ref(2)P (1:1000, gift from Tamaki Yano at Tohoku Univ.). The following secondary goat antibodies were used at 1:200: anti-rabbit Alexa 488 highly cross-adsorbed (ThermoFisher; A11034), anti-rabbit Alexa 555 cross-adsorbed (ThermoFisher; A21429), anti-mouse Alexa 488 highly cross-adsorbed (ThermoFisher; A11029), anti-mouse Alexa 555 highly cross -absorbed (ThermoFisher; A21424) and anti-rat Alexa 647 (Jackson; 112-606-143).

### Quantification and Statistical Analysis

To quantify the distribution of ER and HAGox in the olf hair cells, a single optical section of Airyscan image was selected. Using the PlotProfile function of ImageJ/ Fiji, intensity profiles of a line across the middle of the hair were obtained for 5-7 individual hairs. Normalized and averaged values were calculated by Python.

For envelope curvature, TEM images were analyzed by Kappa in the FIji plugin. Violin plots were created using the median value extracted from the point curvature of each envelope fragment obtained in the analysis. The Mann-Whitney u-test was used for statistical analysis.

### Cell culture experiment

*Drosophila* Schneider 2 cells were cultured in Sf-900II SFM serum-free medium (ThermoFisher/Gibco) supplemented with Penicillin and Streptomycin. Cells were seeded in the 6-well plate (Iwaki). Expression vectors (pUAST-based vectors and pWA-Gal4 (Gal4 linked to the actin 5C promoter) using TransIT-Insect Transfection Reagent (Takara) according to the manufacturer’s protocol. Forty-eight hours later, cells were reseeded onto the Con-A coated cover glasses (15 mm) on a Parafilm in a humid chamber for 30 to 60 min. Coverslips were covered with 0.5 mg/ml concanavalin A (Sigma-Aldrich) in water and air-dried (Rogers et al., 2002). Cells on the cover glass were washed three times and fixed with 4% paraformaldehyde in PBS, blocked in 0.5% BSA, 0.1% Triton X-100 in PBS, and stained with the set of antibodies listed in Supplement Table 1. FlexAble CoraLite® Plus 488 Antibody Labeling Kit for Mouse IgG1 was used to label the mouse anti-Cnx antibody for double labeling with two mouse antibodies. Samples were mounted in VECTASHIELD Mounting Medium (Vector Laboratories), SlowFade Diamond Antifade Mountant (without DAPI) or ProLong Glass Antifade Mountant (without DAPI, ThermoFisher), and observed with an Olympus FV1000 confocal microscope. Image deconvolution was performed by Richardson–Lucy algorithm implemented to ImageJ.

### Immunoprecipitation and Western blotting

Transfected cells were harvested from the culture dishes by pipetting, washed two times with PBS, and lysed in 200 µl of RIPA buffer (50mM Tris-HCl pH7.5, 150mM NaCl, 1% NP40, 0.5% Deoxycholate, 0.1% SDS, supplemented with protease inhibitor cocktail) per 35mm well. Immunoprecipitation was performed with antibodies conjugated to magnetic beads. Western blotting was performed by electrophoresis in pre-casted acrylamide gradient gel (SuperSep Ace, Fuji Film), transferred to PVDF membrane (iBlot2 Transfer Stacks, PVDF), incubated with antibody solutions with iBind Automated Western System (Thermo Fisher Scientific). The signal was detected with chemiluminescence (ECL™ Prime Reagent and Davinch-Chemisystem imager).

### Sample preparation for mass spectrometry

#### Experiment 1

S2 cells transfected with pUAST-attB HA-gox-Flag were lysed in RIPA buffer and immunoprecipitated with magnetic beads coated with anti-HA or anti-Flag antibody. Immunoprecipitates were electrophoresed in SDS-polyacrylamide gel and stained with Silver Stain kit (Fujifilm). Each lane was cut into 16 fragments and analyzed by mass spectroscopy in RIKEN BDR Proteomics Core facility. Proteins enriched by immunoprecipitation with anti-Flag (cytoplasmic site) but not in anti-HA (ER lumen) were identified as candidate Gox interactors.

#### Experiment 2

S2 cells co-transfected with pUAST-attB gox-Flag and pUAST-attB Ref(2)PMyc were lysed in RIPA buffer and immunoprecipitated with magnetic beads coated with anti-Myc, anti-Flag or anti-HA (negative control) antibody. Whole lanes of Silver-stained, short-electrophoresed gel were analyzed.

#### Experiment 3

Anti-Flag immunoprecipitate of experiment 2 was electrophoresed, and the molecular mass range of 25k to 37k was cut and analyzed for ubiquitinated peptides.

Gel fragments were reduced with dithiothreitol and alkylated with iodoacetamide. Digestion was carried out using MS-grade trypsin (Thermo Fisher Scientific). For ubiquitination site identification, additional digestion with pepsin (Promega) was performed. Resultant peptides were extracted using 1% trifluoroacetic acid and 50% acetonitrile solution, dried under vacuum, and reconstituted in 2% acetonitrile and 0.1% trifluoroacetic acid solution.

### LC-MS/MS analysis

In the Experiment 1, mass spectra were acquired using an LTQ-Orbitrap Velos Pro (Thermo Fisher Scientific) coupled to a nanoflow UHPLC system (ADVANCE UHPLC; AMR Inc.) with an Advanced Captive Spray SOURCE (AMR Inc.). Peptide mixtures were loaded onto a C18 trap column (CERI, ID 0.1 mm × 20 mm, particle size 5 μm) and fractionated using a C18 L-column (CERI, ID 0.075 mm × 150 mm, particle size 3 μm). A linear gradient from 5% to 35% solvent B over 20 minutes at 300 nL/min flow rate was used. Solvent compositions were 100% H_2_O with 0.1% formic acid (Buffer A) and 100% acetonitrile with 0.1% formic acid (Buffer B). The mass spectrometer operated in data-dependent mode with thirteen successive scans. Initially, full-scan MS was conducted at 60,000 resolution over the range of 350–2000 m/z using an orbitrap. MS/MS scans were performed on the twelve most intense ion signals using collision-induced dissociation (CID) with a normalized collision energy of 35%, 2 m/z isolation width, and 90-second activation time.

In the Experiment 2 and 3, Mass spectra were acquired using an Orbitrap Eclipse (Thermo Fisher Scientific) coupled to a nanoflow UHPLC system (Vanquish; Thermo Fisher Scientific). Peptide mixtures were loaded onto a C18 trap column (PepMap Neo Trap Cartridge, ID 0.3 mm × 5 mm, particle size 5 μm) and separated on a C18 analytical column (Aurora, ID 0.075 × 250 mm, particle size 1.7 μm, IonOpticks). The peptides were eluted at a flow rate of 300 nl/min using the following gradient: 0% to 2% solvent B over 1 minute, 2% to 5% over 2 minutes, 5% to 16% over 19.5 minutes, 16% to 25% over 10 minutes, 25% to 35% over 4.5 minutes, a sharp increase to 95% over 4 minutes, hold at 95% for 5 minutes, and finally re-equilibration at 5%. The Orbitrap operated in a data-dependent mode with a 3-second cycle time. full-scan MS was collected at 60,000 resolution, the mass range 375 -1500 m/z, using a standard AGC and maximum injection time of 50 ms. MS/MS scan was triggered from precursors with intensity above 20,000 and charge states 2-7. Quadrupole isolation width was 1.6 m/z, with normalized HCD energy of 30%, and resulting fragment ions recorded in Orbitrap at 15,000 resolution with standard AGC target and maximum injection time of 22 ms. Dynamic exclusion was set to 20 seconds.

### Data processing

The raw data files were searched against the drosophila melanogaster dataset (Uniprot Proteome UP000000 803) with the common Repository of Adventitious Proteins (cRAP, ftp://ftp.thegpm.org/fasta/cRAP) for contaminant protein identification, using Proteome Discoverer 2.5 software (Thermo Fisher Scientific) with MASCOT ver.2.8 search engine, with a false discovery rate (FDR) set at 0.01. The number of missed cleavages sites was set as 2. Carbamidomethylation of cysteine was specified as a fixed modification, while oxidation of methionine and acetylation of the protein N-terminus were treated as variable modifications. To identify ubiquitination sites, di-glycine of lysine, and deamidation of glutamine and asparagine were included as additional variable modifications.

### Data availability

Raw image data and uncut images of Western blot are available in SSBD repository (https://ssbd.riken.jp/repository/375/). All mass spectrometry data have been deposited to ProteomeXchange Consortium via jPOST with the accession number PXD053779 and JPST003205 (Preview URL https://repository.jpostdb.org/pre-view/78763230166c42ea45393c, Access key 9275).

## Acknowledgment

We thank the Kyoto *Drosophila* Stock Center, National Institute of Genetics, Bloomington *Drosophila* Stock Center, Vienna *Drosophila* Resource Center, The Drosophila Genetic Resource Center and the Developmental Studies Hybridoma Bank, Tamaki Yano for providing fly stocks and antibodies. We thank assistance from Mai Shibata for cell culture experiments, Hikari Mori for image segmentation, Kenta Onoue and Satoko Okayama for electron microscopy experiments. We thank Noboru Mizushima, Tetsuya Takeda and Koji Takei for discussion, Takefumi Knodo and Fubito Natatsu for comments on the manuscript, Douglas Sipp for editing and members of the Hayashi lab for their comments on the manuscript.

## Author contributions

S.H. conceived the study. S.I. performed the fly experiments, electron microscopy, and image data segmentation. H.W. performed cell culture experiments and protein interaction assays. Y.I. analyzed ECM distribution in UAS-shi^ts^ flies. R.N. performed mass spectrometric analyses. T.I. A.I. L.C. K.M. performed FIB-SEM data acquisition and assisted 3D structure analysis. S.H. and S.I. analyzed the data and wrote the manuscript with input from all of the authors. All of the authors discussed the results and commented on the manuscript.

## Additional Information

Supplementary Information is available for this paper. Reprints and permissions information is available at www.nature.com/reprints.

## Materials & Correspondence

Correspondence and requests for materials should be addressed to Shigeo Hayashi.

---

**Extended Data Figure 1.**
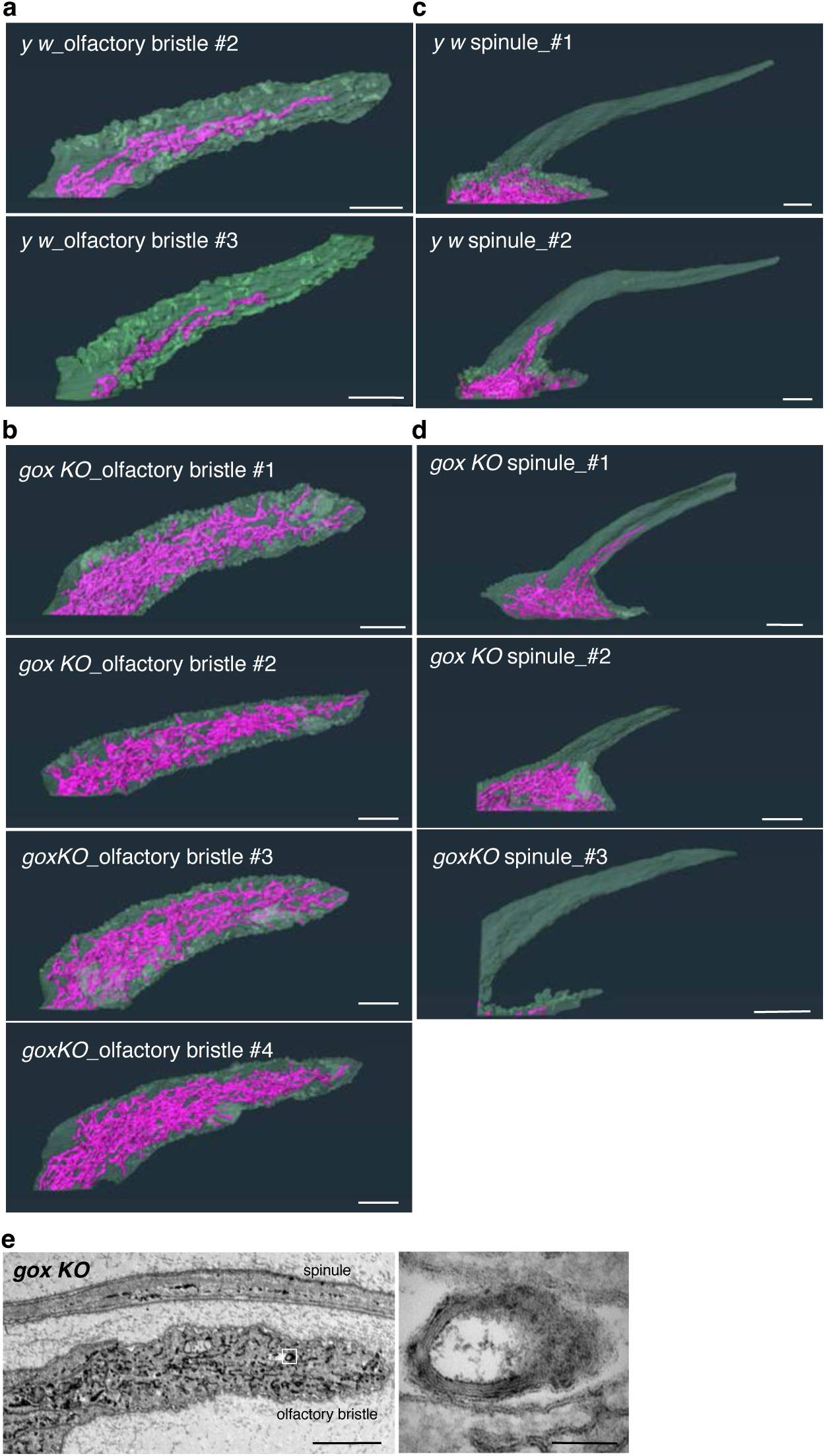
FIB-SEM reconstructions of ER and plasma membrane in multiple hair cells of olf and spinule sampled from third antennal segment at 42h APF. **a,** Control olf. **b,** Control spinule. **c,** *gox*^1^ olf. **d,** *gox^1^*spinule. **e,** TEM view of the *gox* mutant olf. ER whorl structure is enlarged. Bar: 1µm.

**Extended Data Figure 2.**
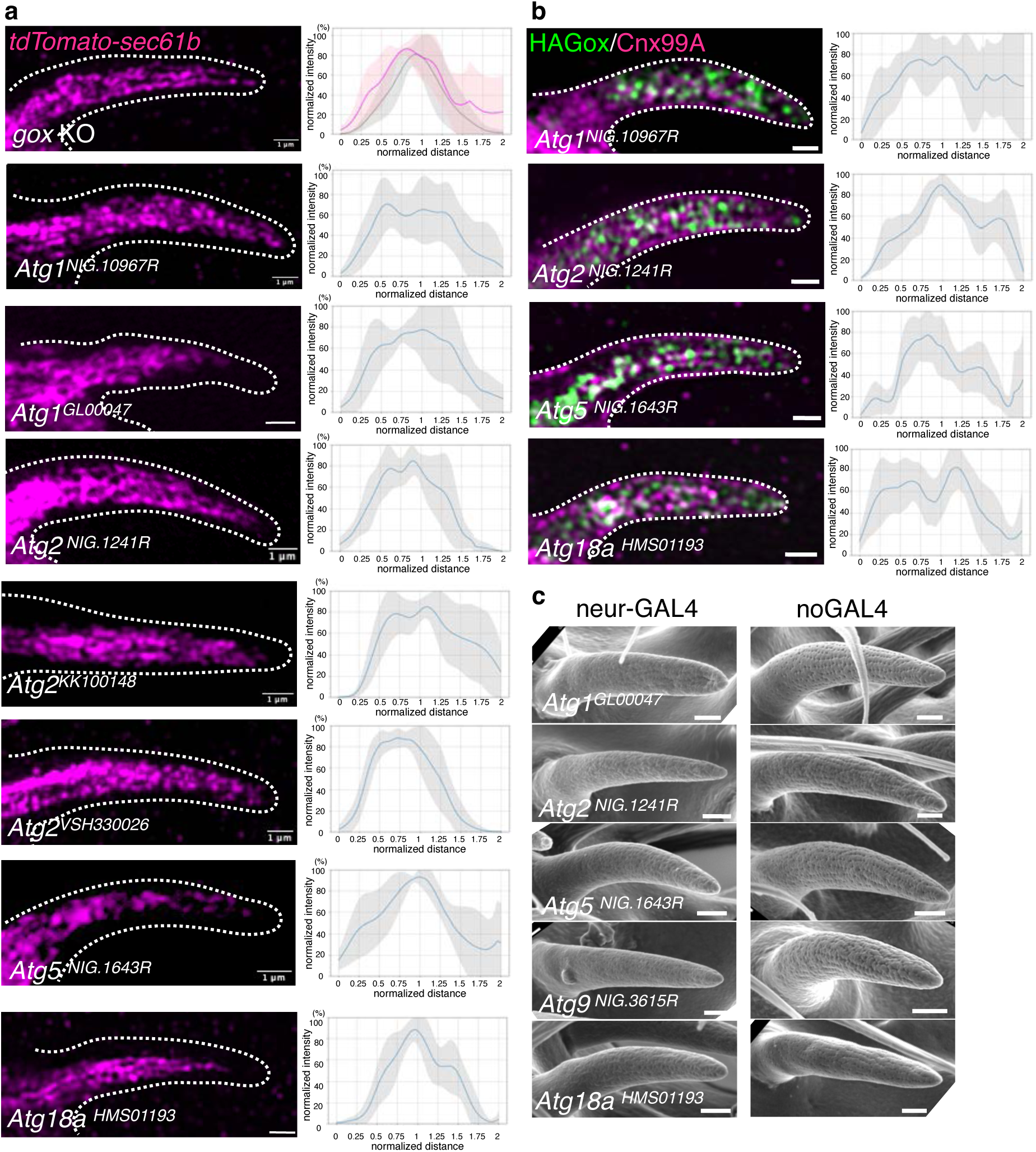
**a,** ER distribution in *gox* mutants, two ATG1 RNAi, and ATG2 RNAi. **b**, HA-Gox distribution in ATG1, ATG2, ATG5 and ATG18a RNAi showing the reduction of subcortical Gox. **c**, Olfactory hairs of ATG RNAi experiments. All showed reduced nanopores compared to the control. Bar: 1µm.

**Extended Data Figure 3.**
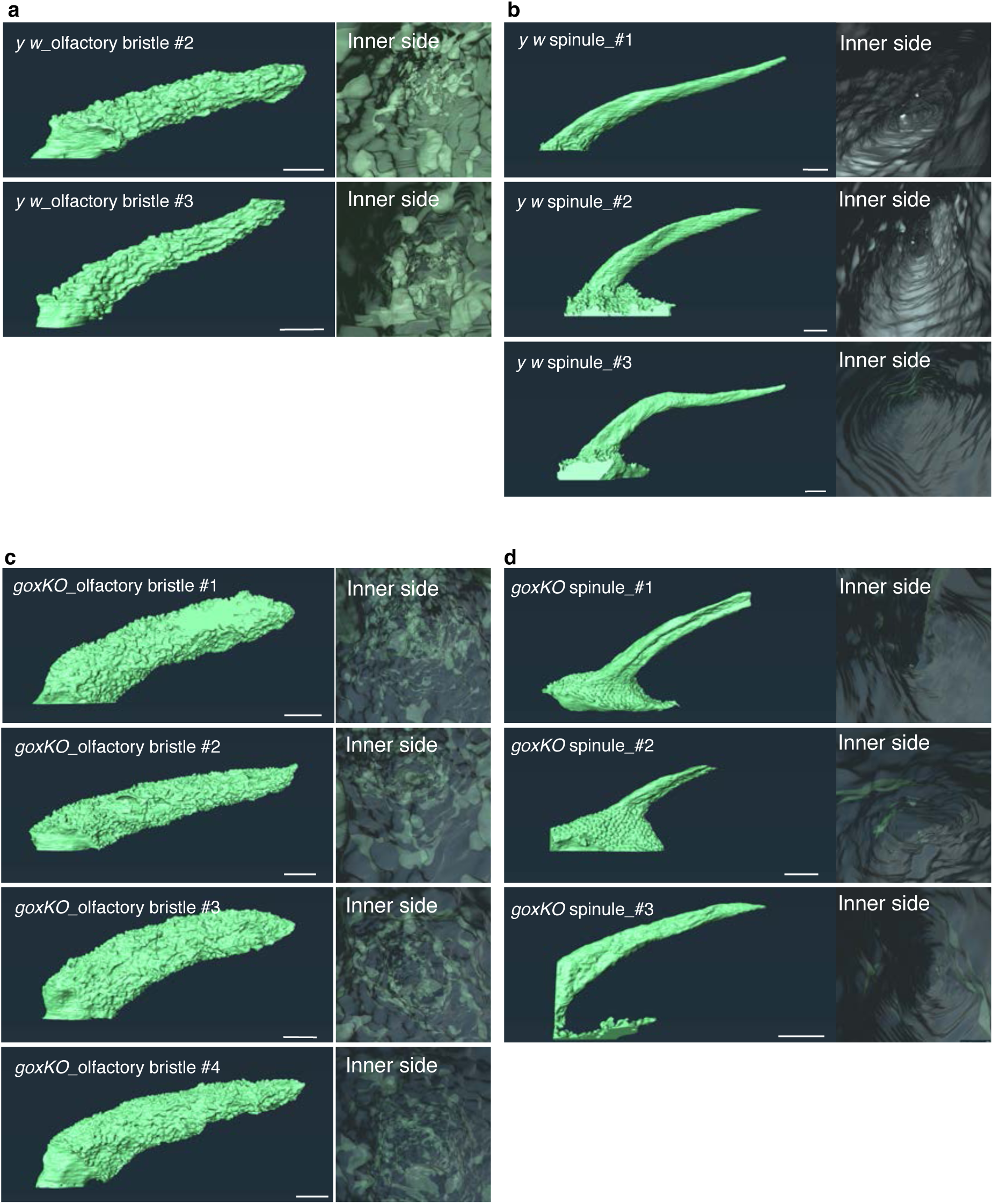
FIB-SEM analysis of plasma membrane. External and internal views of **a,** *y w* (control) olf hair cells. **b**, *y w* spinule, **c,** *gox* mutant olf and **d,** *gox* mutant spinule, are shown.

**Extended Data Figure 4.**
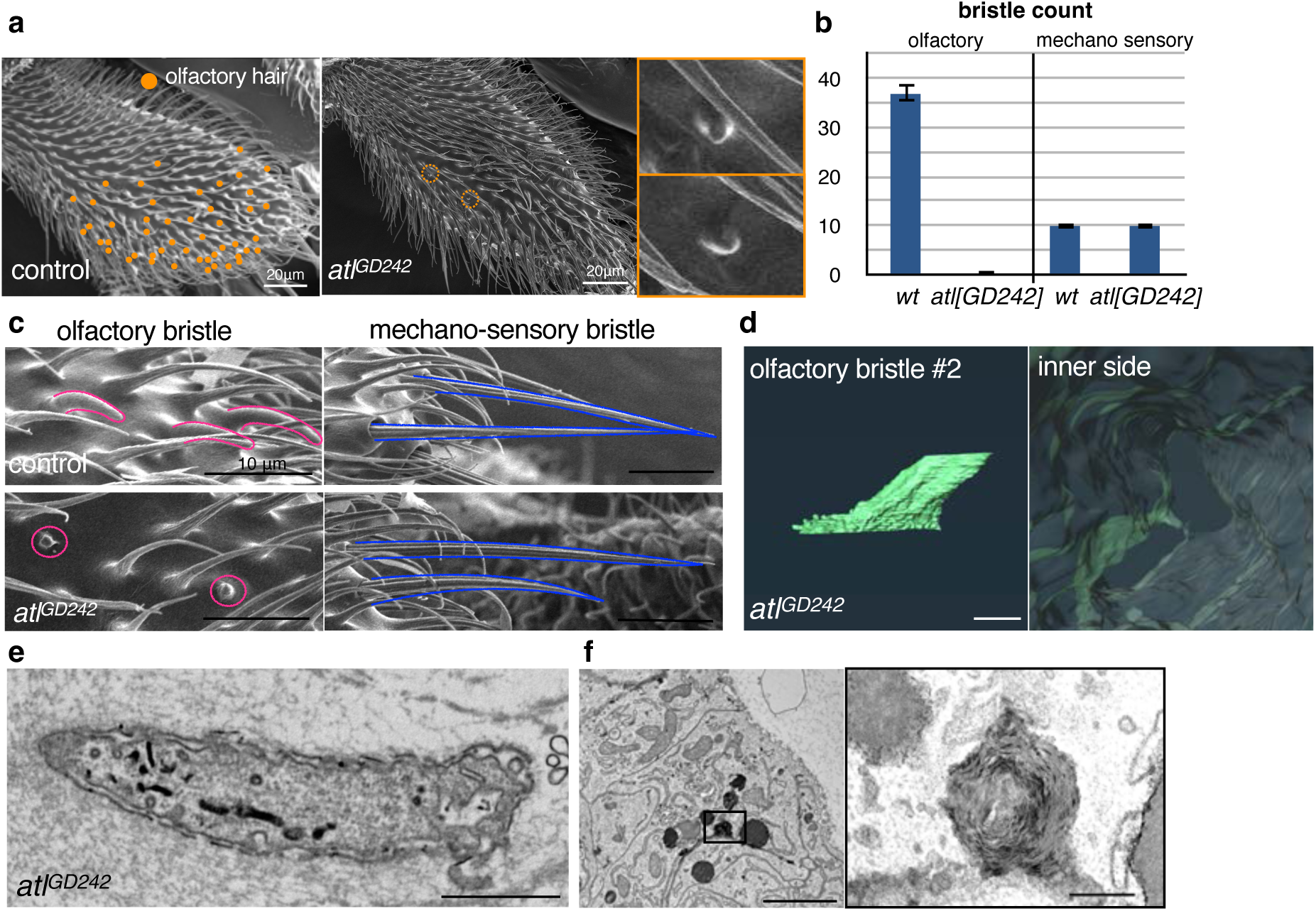
Atlastin RNAi phenotype. **a,** SEM views of maxillary palp of *y w* (control) and neur>atl RNAi flies. **b**, Count of olf and mech bristles. N of palp = 18 (wt), 16 (atl RNAi). **c,** Enlarged views of olf and mech bristles. Note that *neur-Gal4* used in this experiment is active in both bristles. **d,** FIB-SEM view of plasma membrane trace of *atl* RNAi olf hair cells. **e,** Longitudinal TEM section of *atl* RNAi olf hair cells. Note straightened envelopes. **f,** TEM view of condensed ER in the basal region of olf hair cell. Enlargement shows a multi-membrane “whorl” phenotype.

**Extended Data Figure 5.**
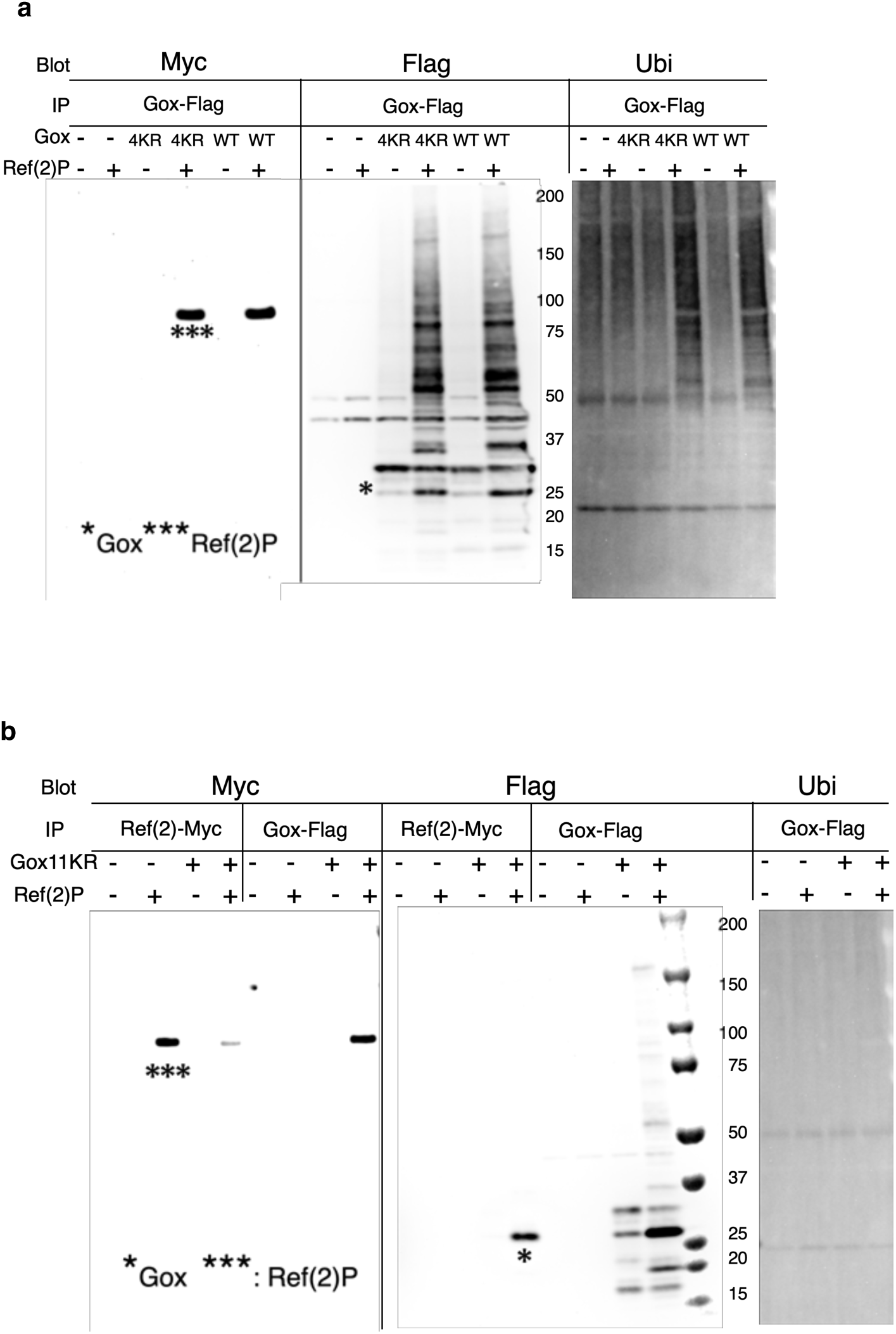
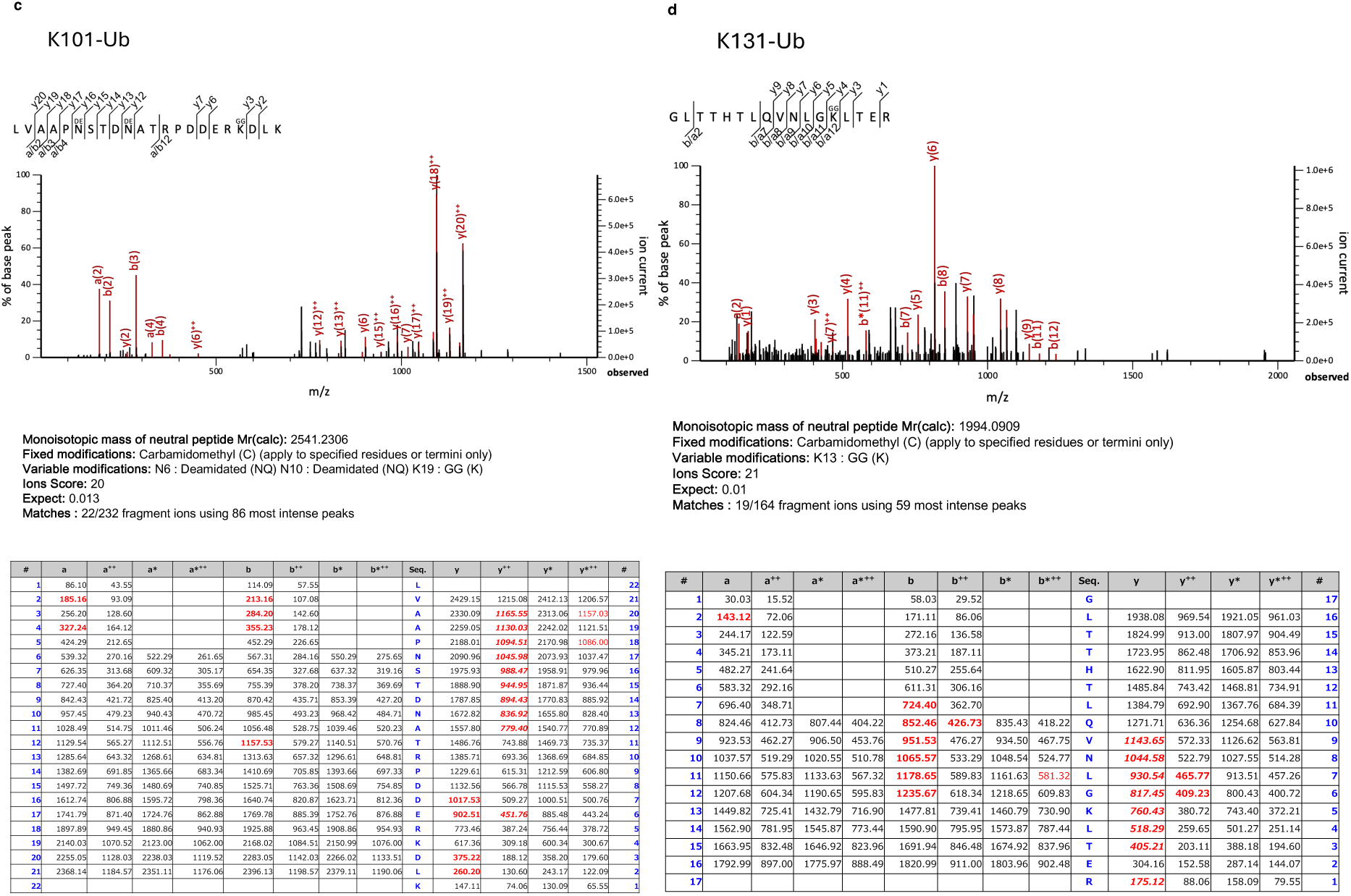
Analysis of Gox ubiquitination. **a** Gox4KR mutant and GoxWT were both bound to Ref(2)P and present as high-molecular-weight ubiquitinated forms. **b**, Property of Gox11KR mutant. Ref(2)P associated with Gox11KR, and increased its amount. But caused no change in ubiquitination. The phenotype was identical to Gox8KR. **c, d**, Identification of ubiquitination sites at K101 and K131. di-Glycilated lysine was identified at those sites.

**Extended Data Figure 6.**
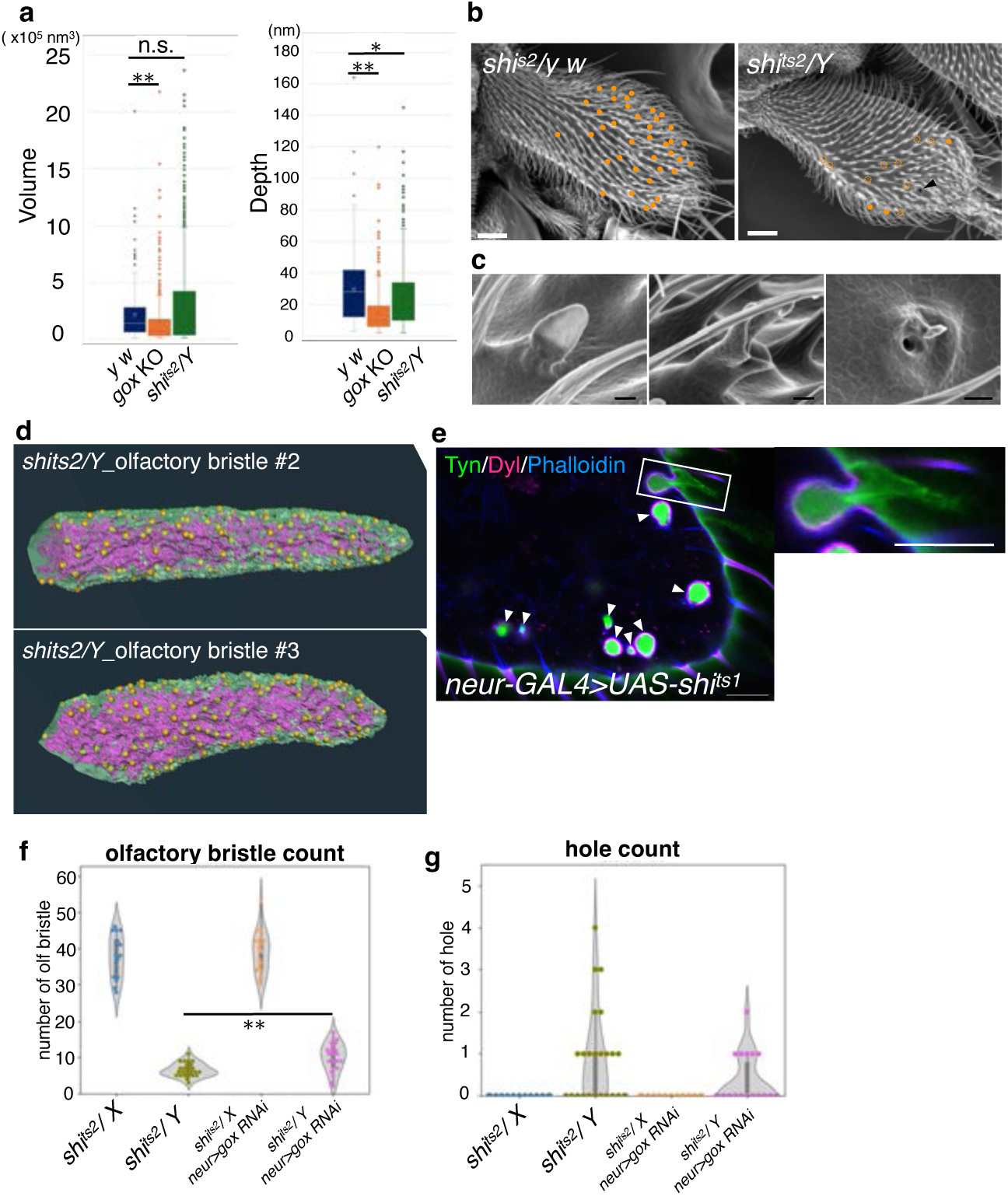
Analysis of Dynamin-deficient olf hair cells. **a,** PMI volume and depth measured from FIBSEM reconstructions of three olf hair cells of *y w, gox* and *shi^ts^*^2^*/Y* pupae. **b**, Phenotype of maxillary palp bristles. Female (*sh^ts^*^2^*/X,* control) and male *(sh^ts^*^2^*/Y,* experimental) progenies from the same batch of heat-treated flies are compared. Filled dot and dotted circle indicate olf bristles of normal length and abnormal morphology, respectively. **c**, Enlargement of abnormal olf bristles of *sh^ts^*^2^*/Y*. Left to right: short, branched and hole phenotype. **d**, FIB-SEM views of additional olf hair cells of *shi^ts^*^2^*/Y* showing ER, plasma membrane, and their contact site (yellow dot). **e**, Maxillary palp expressing *UAS-shi^ts^*^1^ driven by the *neur-Gal4* driver. Apical ECM marker: mVenus-Tyn and mScarlet-Dyl. Arrowhead indicates the lumen of invaginated olf cells. Enlargement shows a hair cell-like remnant of Tyn left outside of the invaginated olf cell. **f, g,** Count of olf bristle (**f**) and hole phenotype (**g**) per maxillary palp. Note that *neur>gox RNAi* partially suppressed the *sh^ts^*^2^*/Y* phenotype. Bar: 20µm (b), 1 µm (c, d). Asterisk: **, p<0.001 (**a, f**); *, p<0.05 (**g**). Statistical significance was determined by Mann-Whitney U test.

## Supplementary Information

**Supplementary Table 1.** Reagent List.

**Supplementary Table 2.** List of proteins identified in the mass spectrometry analyses.

**Supplementary Table 3.** Primer List.

**Supplementary Movie 1.** 3D view of the plasma membrane of the olf hair cell.

